# Interindividual variability in flower pickiness by foraging bumblebees

**DOI:** 10.1101/2024.12.19.629386

**Authors:** Mélissa Armand, Christoph Beckenbauer, Aurore Avarguès-Weber, Mathieu Lihoreau, Tomer J. Czaczkes

## Abstract

Pollinators navigate complex and heterogeneous “flower markets”, where floral resources vary in quality, availability, and spatial distribution. Bumblebees, as generalist foragers, visit numerous flowers during their foraging bouts, yet the factors influencing their flower choices and the individual differences in foraging behaviour remain poorly understood. Here, we tested how bees adjust their foraging in response to different reward structures. *Bombus terrestris* workers completed three foraging bouts in two artificial flower environments: one simulating a favourable environment with patches alternating high- and low-quality flowers (40% vs. 20% w/w sucrose solution), and the other a challenging environment with high-quality flowers alongside unrewarded ones (40% w/w sucrose solution vs. plain water). We hypothesised that bees would improve their foraging efficiency in both environments, but more rapidly in the more extreme one, where the greater reward difference creates stronger pressure to learn quickly. In both conditions, bees increased their sucrose intake per unit time over bouts. We also observed consistent differences in flower selectivity among individuals: in the favourable environment, bees that first visited high-quality flowers focused on them and avoided low-quality ones (became “picky”), while bees that first visited low-quality flowers kept visiting both types. Despite these differences, bees across environments and pickiness levels all reached similar sucrose intake rates by the third foraging bout, either by becoming more selective, collecting more sucrose solution, or reducing time spent foraging. These findings highlight the adaptability of bee foraging and suggest that early flower experiences may contribute to lasting individual differences in foraging behaviour.

**Significance statement:** Bumblebees are highly efficient pollinators of wild plants and commercial crops, yet their foraging behaviour varies notably among individuals. Understanding why bees differ in their foraging decisions is crucial, as it affects how they collect resources. We show that individual *Bombus terrestris* workers rapidly adapt to flower patches with different reward qualities. Remarkably, a bee’s first flower visits have lasting effects: individuals that started with high-quality flowers consistently favoured them and avoided low-quality ones, while those that started with low-quality flowers continued to visit both types over time. Despite these differences, bees reached similar nectar intake rates after just three foraging trips, regardless of how selective they were. Our findings show that bees can quickly adjust their foraging behaviour, and that early experiences play a key role in how they exploit resources across different environments.

## Introduction

Pollinators such as bees, bats, and birds forage in diverse and dynamic “flower markets”, where collecting food efficiently is a complex task. Optimal Foraging Theory (OFT) posits that animals forage in ways that optimise net energy gain, i.e., by maximising energy intake per unit of time (Pyke, 1984; Schmid-Hempel et al., 1985; Pyke and Starr, 2021). In heterogeneous environments, there is no single way to improve foraging efficiency; animals may forage faster, select higher-quality resources, or collect larger quantities. Optimising foraging may thus be particularly challenging for generalist bees like the central-place foraging bumblebees, which can exploit hundreds of flowers each day with rates of 10 to 20 flower visits per minute (Heinrich, 1979a). Yet, bees vary considerably in how they exploit floral resources (Woodgate et al., 2016). Despite extensive research on bee foraging (see review by Sommer et al., 2022), the causes of individual differences in foraging behaviour remain unclear. Although environmental variability likely affects bee foraging, how it contributes to consistent differences between individuals is not well understood.

The natural foraging environments of bees are highly dynamic, with nectar availability and quality fluctuating across flowers and throughout the day (Goulson, 2003). For example, aging flowers can change colour and reduce nectar rewards (Weiss, 1991; Willmer, 2011). Many flowers lack nectar entirely, with up to 70% on a single plant being nectarless (Real and Rathcke, 1988; Cresswell, 1990; Thakar et al., 2003). Nectar quality also varies widely, with sugar concentrations ranging from 10% to 80% w/w (Kevan and Baker, 1983). In addition to environmental variability, bees face internal constraints that influence their foraging decisions. Wing wear, for instance, can lead bumblebees to favour denser floral patches to minimise flight effort (Foster and Cartar, 2011). Body size also plays a role: larger bees carry more nectar and tend to forage more efficiently (Heinrich, 1979a; Goulson et al., 2002; Spaethe and Weidenmüller, 2002). Experience shapes foraging as well, with experienced bees making faster, more direct flights (Osborne et al., 2013; Woodgate et al., 2016), although learning ability varies among individuals (Evans and Raine, 2014). Given the high energetic cost of foraging (Darveau et al., 2014; Vaudo et al., 2016), bees must balance environmental variability and internal constraints to optimise efficiency.

Foraging bees typically collect nectar from multiple flower types over their lifetimes (Hagbery and Nieh, 2012; Russell et al., 2017), and their floral choices vary across environments and individuals. Naïve bees often show innate preferences for violet or blue flowers (Gumbert, 2000; Raine and Chittka, 2007), larger flowers that are easier to locate (Ohara and Higashi, 1994; Spaethe et al., 2001), and dense flower patches (Makino and Sakai, 2007). Bumblebees also exhibit flower constancy, the tendency to forage predominantly on a single flower species (Chittka et al., 1999), though they do so more flexibly than honeybees (Osborne and Williams, 2001; Gegear and Laverty, 2004). However, both innate preferences and constancy can be overridden with experience (Gumbert, 2000; Makino and Sakai, 2007). While bees generally prefer higher sugar concentrations (Whitney et al., 2008), they do not always choose nectar-rich flowers. For instance, Abrol (2006) showed that honeybees favoured plants with high flowering density and lower nectar concentration over patches with fewer, nectar-rich flowers. Similarly, smaller-size bumblebees invested equal effort in memorising flowers with high and low sugar concentration (Frasnelli et al., 2021). Preferences for nectar volume have received less attention, and findings are inconsistent (Menzel and Erber, 1972; Waddington and Gottlieb, 1990; Lihoreau et al., 2011), although bees appeared more responsive to changes in nectar concentration than volume (Cnaani et al., 2006). Floral choices are also shaped by bees’ energetic demands, as they adjust foraging to meet both individual needs and those of the colony (Molet et al., 2008; Vaudo et al., 2016; Hendriksma et al., 2019).

Depending on environmental conditions, bees may use different foraging “tactics” to navigate floral resources. In simpler flower patches, they often follow a nearest-neighbour rule, moving sequentially to the closest unvisited flower until resources are depleted (Saleh and Chittka, 2007; Ohashi et al., 2007). In denser or more aggregated patches, bees adopt a near-far search rule, making short flights between rewarding flowers and staying longer in nectar-rich areas, but leaving quickly or taking longer flights when rewards decline (Dukas and Real, 1993; Chittka et al., 1997; Cartar, 2004). At larger spatial scales, bees alternate between exploration and exploitation flights to establish foraging routes (Kembro et al., 2019). A common pattern is traplining: repeatedly visiting flowers in a fixed sequence to maximise rewards while avoiding already depleted sites (Ohashi et al., 2007; Lihoreau et al., 2010). Bees gradually refine these routes to balance shorter paths with prioritising high-reward flowers (Reynolds et al., 2013). Still, foraging patterns in heterogeneous environments are highly variable (Lihoreau et al., 2012), and individual bees differ in patch fidelity, the exploration–exploitation balance, and the duration and frequency of foraging bouts (Woodgate et al., 2016). We propose that environmental conditions shape lasting individual differences in flower choice and foraging behaviour.

Here, we conducted a lab-based experiment to test how bees improve foraging efficiency in heterogeneous flower environments. We examined how the quality difference between good and poor flowers within patches influences the foraging behaviour of *Bombus terrestris* workers. Individual bees completed three consecutive foraging bouts in two distinct artificial flower environments. The first simulated a favourable environment, with patches containing alternating high- and low-quality flowers (“High vs. Low” environment; 40% vs. 20% w/w sucrose solution). The second represented a challenging environment, with patches of rewarding flowers alongside unrewarded ones (“High vs. Water” environment; 40% w/w sucrose solution vs. plain water). We hypothesised that environmental context would shape bees’ subsequent foraging behaviour. Specifically, we predicted that the more extreme conditions in the High vs. Water environment would allow bees to learn flower quality more rapidly and improve foraging efficiency faster than in the High vs. Low environment. Nonetheless, we expected that bees in both environments would similarly increase their energy intake rate with experience, consistent with optimal foraging theory (Pyke, 1984; Schmid-Hempel et al., 1985).

## Material and Methods

### Bees

Six commercial *B. terrestris* colonies (Koppert, The Netherlands) were maintained under controlled laboratory conditions (22–24°C, 14:10 light:dark cycle; lights on from 6:30 a.m. to 8:30 p.m.) with ad libitum pollen. Colonies were housed in wooden nestboxes connected to individual flight arenas (60 x 50 x 35 cm) via transparent tubes fitted with shutters (see apparatus in **Fig. 1**). Between experimental sessions (i.e., pre-training), bees had unrestricted access to six artificial flowers providing ad libitum 20% w/w sucrose solution through a damp cotton mesh soaked in solution and protruding through a central hole. Bees that foraged regularly were individually marked on the thorax with uniquely coloured, numbered tags and selected for the experiment.

**Figure 1:**
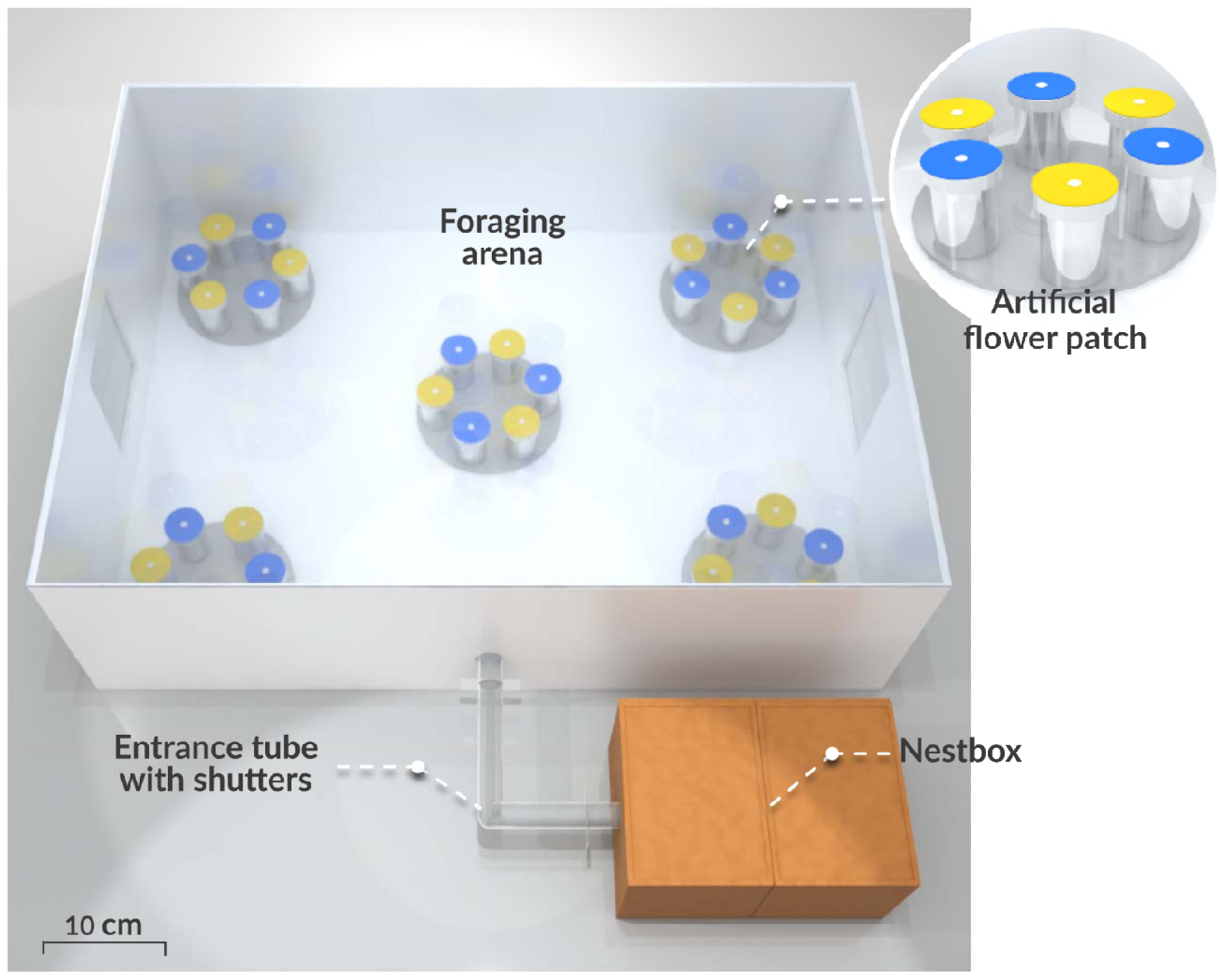
Top view of the scaled experimental setup. The 3D model, created in Blender, illustrates the layout of pattern A of the flower environment (see **Supplement S5** for all layout patterns). Flower patches were positioned 15–50 cm from the arena entrance and mounted on transparent white circular plastic bases (15 cm diameter, 0.5 cm thick), with flowers within each patch spaced 1.5 cm apart. Flowers were removed and rearranged after each foraging bout using the arena’s side doors.

A total of 52 bees from four colonies were tested over 20 days (October–November 2021; see **Supplement S1** for colony details). Additionally, six bees from two other colonies were tested in January 2024 in a complementary experiment to determine whether individuals collected the entire sucrose solution drop from the artificial flowers or only a portion of it. We conducted this control to validate whether sucrose solution consumption could be accurately estimated based on reward collection. This setup mirrored the main experiment but reversed flower colours and included weighing flowers before and after each foraging bout. Results showed that bees typically either fully collected the sucrose solution or rejected the flower without drinking (see **Supplement S2** for detailed analysis).

### Artificial flowers

Each flower consisted of a transparent plastic cup (38 mm diameter, 70 mm height) topped with a coloured laminated paper disk of the same diameter. Flowers used during the experiment were either blue (peak reflectance ≈ 450 nm) or yellow (peak reflectance ≈ 530–700 nm; see **Supplement S3** for reflectance curves). Sucrose solution was pipetted onto a small white dot in the centre of each flower to aid the bees in locating the rewards (Heuschen et al., 2005; see **Fig. 1**). During pre-training, flowers were made of two halves, one blue and one yellow, following the method of Raine and Chittk (2008). These bicoloured flowers ensured bees associated sucrose rewards equally with both colours, which were later used as distinct flower types in the experiment.

### Experiment

Individual bees completed three consecutive foraging bouts on a flower environment consisting of five dimorphic flower patches, each with six flowers arranged in a circle of alternating blue and yellow colours. This ensured that neighbouring flowers were always of different colours. Patches were arranged using four distinct layout patterns (see **Supplement S5**), and each bee experienced a random sequence of three different patterns. This pseudo-random design prevented reliance on spatial memory while keeping layouts consistent across bees.

In the main experiment, yellow flowers consistently offered high-concentration sucrose solution to counteract bees’ innate preference for blue (Gumbert, 2000) and avoid a colour bias. To further test the robustness of the setup, colour assignments were reversed in the complementary experiment. Bees were tested on one of two flower environment types:

1. **High vs. Low flower environment** (N = 37 bees from 6 colonies): For 31 bees, yellow flowers contained 14 μL of concentrated sucrose solution (40% w/w) and blue flowers 14 μL of diluted sucrose solution (20% w/w). For the six bees in the complementary experiment, the colours were reversed. This simulated a favourable environment where all flowers were rewarding but varied in quality. We expected bees in this environment to be less selective in their flower choices and to show slower improvement in foraging efficiency compared to the environment with more extreme differences in flower rewards described below.
2. **High vs. Water flower environment** (N = 15 bees from 3 colonies): Yellow flowers contained 14 μL of concentrated sucrose solution (40% w/w), and blue flowers 14 μL of plain water (tap water). This represented a more challenging environment, with only half of the flowers offering a reward. We expected bees in this environment to be highly selective and show rapid improvement in foraging efficiency.

In both environments, each flower type provided a total of 210 μL of sucrose solution per bout, enough for a bee to fill its crop by visiting a single flower type. This volume exceeds the average crop capacity of bumblebee foragers (120–180 μL; Lihoreau et al., 2010). To prevent chemical marks from influencing flower choices, all flower surfaces were wiped clean with a paper towel moistened with 70% ethanol between foraging bouts (Goulson et al., 2000; Saleh et al., 2007).

After the third training bout, bees completed a final test bout in which the arena contained two monochromatic flower patches, one yellow and one blue, both unrewarded (plain water). The patches were positioned against the back wall, facing the arena entrance, and the bee’s first flower visit was recorded as an indicator of its flower type preference.

### Data collection and analysis

Each foraging bout was video-recorded using a camera (Sony HDR CX405) mounted above the flight arena. Bee behaviour was analysed using an event-logging software (BORIS, v8.6), recording: (1) foraging bout duration (i.e., time from entering the arena to returning to the nest), (2) flower visits (bee landing on a flower), (3) flower revisits (bee landing on a previously visited flower within the same bout), and (4) reward collection (bee licking a drop of sucrose solution for more than five seconds). Based on video analysis, bees were classified as either rejecting the flower or accepting and fully consuming the sucrose drop (see **Supplement S6** for video examples). This categorisation was supported by the complementary experiment, in which flowers were weighed before and after each foraging bout (see **Supplement S2** for methodology and results). Bees that accepted a flower overwhelmingly consumed the entire drop, regardless of whether it was high- or low-concentration sucrose.

Data processing was performed with Python (v3.11, Python Software Foundation, 2023) using the following libraries: *pandas* (McKinney, 2010) for data structuring, *seaborn* (Waskom, 2021) and *Matplotlib* (Hunter, 2007) for data visualisation, and *Scikit-learn* (Pedregosa et al., 2011) for data clustering. Statistical analyses were carried out with R (v4.1, R Core Team 2022) using the packages *glmmTMB* (Brooks et al., 2017) for Generalised Linear Mixed Models (GLMMs) and *emmeans* (Lenth, 2020) for post-hoc comparisons. Model residuals were evaluated using the *DHARMa* package (Hartig, 2020).

Datasets from the main and complementary experiments, where flower colours associated with reward were reversed, were merged for analysis. To account for potential effects of flower colour, experiment type (main vs. complementary) was included as a fixed effect in all models. Individual bees were treated as random effects nested within colonies to account for within-colony variation. All statistical analyses and datasets are available on Zenodo (https://doi.org/10.5281/zenodo.16418595).

### Bee clustering

During initial analysis of bees’ foraging behaviour in the High vs. Low flower environment, we observed substantial dispersion in the proportions of visits to high-quality flowers, collection of high-concentration sucrose, and rejection of low-quality flowers, suggesting notable interindividual variability in flower pickiness. To assess this variation, we applied K-means clustering (k = 2) to the bees based on three behavioural variables measured during the final foraging bout:

1. **Proportion of visits to high-quality flowers**: number of visits to high-quality flowers divided by total visits. A visit is defined as a landing on a flower, regardless of whether the bee collected or rejected the sucrose solution;
2. **Proportion of high-concentration sucrose solution collected**: number of high-quality flowers collected (i.e., solution depleted) divided by total flowers collected;
3. **Proportion of low-concentration sucrose solution flowers rejected**: number of non-empty low-quality flowers visited but not collected (i.e., bees inspected but rejected the sucrose solution), divided by total visits to low-quality flowers.

Each variable was standardised (converted to z-scores by subtracting the mean and dividing by the standard deviation) and clustering was applied directly to these standardised values, identifying two groups of bees with distinct foraging behaviours (Fig. 2B; see detailed analysis in **Supplement S1**):

A. **High pickiness** cluster (N = 28 bees): bees in this group predominantly visited high-quality flowers, collected a high proportion of high-concentration sucrose, and strongly rejected low-concentration flowers
B. **Low pickiness** cluster (N = 9 bees): bees visited high- and low-quality flowers more evenly, collected lower proportions of high-concentration sucrose, and often accepted low-quality flowers instead of rejecting them.

**Figure 2:**
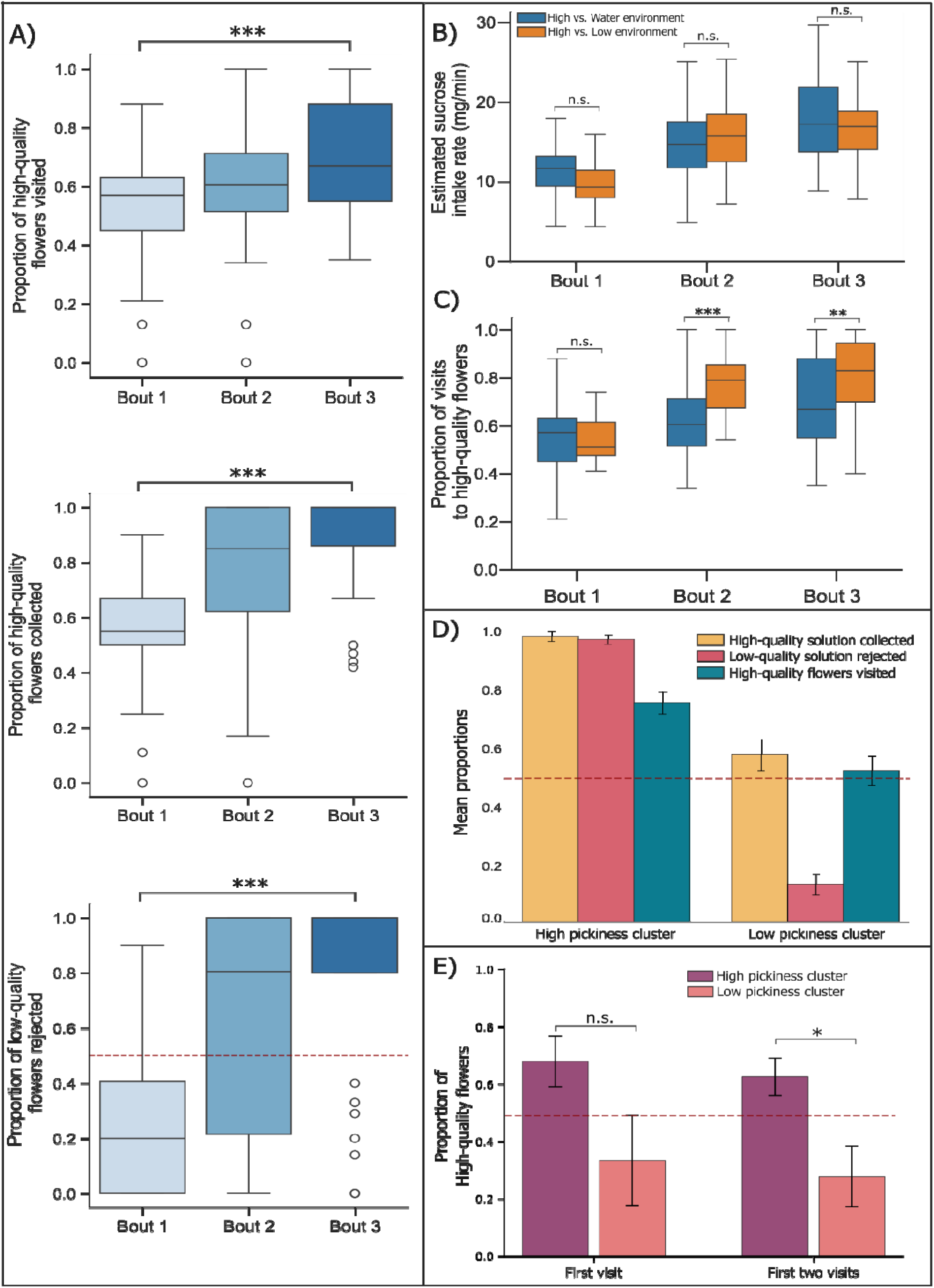
Foraging efficiency across foraging bouts. **A)** Box plots of the proportions of bees (1) visiting high-quality flowers, (2) collecting high-quality flowers, and (3) rejecting low-quality flowers across foraging bouts (N = 37 bees); **B)** Box plots of estimated sucrose intake rates (mg/min) in each flower environment; **C)** Box plots of the proportion of visits to high-quality flowers in each environment. Boxes show the interquartile range (IQR), horizontal lines indicate the median, and whiskers extend to the smallest and largest values within 1.5×IQR. **D)** Bar plots of the mean proportions of variables: (1) visits to high-quality flowers, (2) high-quality flowers collected, and (3) low-quality flowers rejected for each pickiness cluster; **E)** Bar plots of the proportions of high-quality flowers visited during the first flower visits for each bee cluster. Error bars represent standard errors of the mean. Statistical significance is indicated as **\*\*\*** (p < 0.001), **\*\*** (p < 0.01), **\*** (p < 0.05), and **n.s.** (p > 0.05, post-hoc Tukey tests).

### Estimation of absolute sucrose intakes

Absolute sucrose intake per foraging bout was inferred from video analysis, based on the number of high- and low-quality flowers collected and the theoretical sucrose content of 20% and 40% w/w sucrose solutions. These calculations assume that bees consumed the entire sucrose drop upon collection, a behaviour confirmed in our complementary experiment. Calculations were based on the following values (Hofmann, 1977):

1. **20% w/w sucrose solution**: With a density of 1.081 g/mL, a 14 μL drop weighs approximately 15.134[mg, comprising 3.027 mg of sucrose and 12.106 mg of water.
2. **40% w/w sucrose solution**: With a density of 1.176 g/mL, a 14 μL drop weighs approximately 16.464[mg, comprising 6.588 mg of sucrose and 9.882 mg of water.

## Results

We combined data from the main and complementary experiments (the latter reversed colour–reward pairings) for analysis. Despite the reversal, bees showed similar behavioural trends. Naïve bees tended to visit yellow flowers first: 60.9% (18/28) in the main experiment and 67% (4/6) in the complementary experiment. To test whether an innate preference for yellow might have influenced foraging behaviour across bouts, we included “experiment type” (main vs. complementary) as a fixed effect in all models. This variable had no significant effect, indicating that flower colour did not drive the observed behavioural patterns (see **Supplement S2** for model details).

### 1. Foraging efficiency in the favourable vs. challenging environments

We first tested whether environmental context influenced how bees adjusted their foraging behaviour. We predicted that bees in the challenging environment (40% vs. water) would learn to distinguish flower quality faster than those in the favourable environment (40% vs. 20%), and that bees in both environments would improve their foraging efficiency with experience.

#### (A) Estimated sucrose intake rate

Bees in both treatments significantly increased their estimated sucrose intake rate across bouts (GLMM, χ² = 84.23, df = 2, *p* < 0.0001; N = 52 bees), consistent with expectations from Optimal Foraging Theory. Estimated intake rates did not differ between environments at any bout (post-hoc Tukey tests, challenging vs. favourable environment, bout 1: 10.8 mg/min ± 1.19 vs. 12.1 mg/min ± 0.82, z-ratio = 0.93, *p* = 0.35; bout 2: 17.3 ± 1.69 vs. 15.9 ± 1.01, z-ratio = −0.88, *p* = 0.38; bout 3: 18.8 ± 1.81 vs. 19.2 ± 1.17, z-ratio = 0.24, *p* = 0.81; Fig. 2B).

#### (B) Proportion of visits to high-quality flowers

Bees in the challenging environment became more selective over time, visiting high-reward flowers more frequently than those in the favourable environment (GLMM, treatment: χ² = 9.13, df = 1, *p* = 0.0025). There was no difference in bout 1 (post-hoc Tukey test, challenging vs. favourable environment: 58.7% ± 6.0% vs. 52.4% ± 4.4%, t-ratio = −1.18, *p* = 0.24; N = 52 bees), but significant differences emerged in bout 2 (81.5% ± 6.0% vs. 61.0% ± 4.4%, t-ratio = −3.83, *p* < 0.001) and bout 3 (84.1% ± 6.0% vs. 70.1% ± 4.4%, t-ratio = −2.61, *p* = 0.010; Fig. 2C).

### 2. Bees in the favourable environment rapidly increased foraging efficiency

We then examined whether bees foraging on patches with both high- and low-quality flowers favoured the high-concentration sucrose solution or collected solution from both flower types. Over successive foraging bouts, bees significantly increased: (1) their proportion of visits to high-quality flowers, calculated as the number of visits to high-quality flowers divided by total visits (post-hoc Tukey test, bout 1 vs. bout 3: 50.5% ± 4.2% vs. 68.3% ± 4.2%, t-ratio = −6.20, *p* < 0.0001; N = 37 bees; Fig. 2A); (2) their proportion of high-concentration sucrose solution collected, calculated as the number of high-quality flowers collected divided by total flowers collected (post-hoc Tukey test, bout 1 vs. bout 3: 55.6% ± 4.6% vs. 87.7% ± 4.7%, t-ratio = −10.55, *p* < 0.0001; N = 37 bees; Fig. 2A), and (3) their proportion of low-concentration sucrose solution rejected, calculated as the number of non-depleted low-quality flowers visited and not collected, divided by total visits to low-quality flowers (post-hoc Tukey test, bout 1 vs. bout 3: 29.4% ± 7.0% vs. 74.0% ± 6.2%, t-ratio = −5.51, *p* < 0.0001; N = 37 bees; Fig. 2A).

#### (A) Bees varied in their selectivity towards high-quality flowers

The K-means clustering analysis identified two distinct bee clusters: High pickiness (N = 28 bees; 23 from the main experiment and 5 from the complementary experiment) and Low pickiness (N = 9 bees; 8 from the main experiment and 1 from the complementary experiment).

These clusters reflect marked differences in foraging behaviour (Fig. 2D). Bees in the High pickiness cluster visited high-reward flowers in 75% of visits on average, collected 97.8% high-concentration sucrose, and rejected 98.1% of low-quality flowers. In contrast, bees in the Low pickiness cluster visited high-reward flowers in only 51.8% of visits, collected 57.3% high-concentration sucrose, and rejected only 36.6% of low-quality flowers. These proportions illustrate clear behavioural differences between the two groups.

#### (B) Bees’ degree of pickiness was correlated with the quality of the first flowers visited

Bees in the High pickiness cluster tended to have initially visited high-quality flowers more often than those in the Low pickiness cluster (post-hoc Tukey test, High pickiness cluster vs. Low pickiness cluster: 60.1% ± 16.4% vs. 27.6% ± 19.4%; z = 1.48, p = 0.138, N = 37 visits; Fig. 2E). This effect became statistically significant when considering only the first two flower visits (post-hoc Tukey test, High pickiness cluster vs. Low pickiness cluster: 61% ± 8.13% vs. 28.2% ± 11.36%; z = 2.19, p = 0.028, N = 74 visits; Fig. 2E). This finding suggests that early exposure to high-quality flowers may drive bees to become more selective, whereas initial visits to low-quality flowers might reduce selectivity in subsequent choices.

### 3. Bees across environments and clusters improved foraging efficiency via different trade-offs

Next, we examined the foraging behaviour of bees across foraging bouts, comparing flower environment (High vs. Low and High vs. Water) and clusters (High pickiness and Low pickiness). All groups significantly increased their estimated sucrose intake per unit of time with each successive bout (GLMM: χ² = 90.76, df = 2, *p* < 0.0001, N = 52 bees). This improvement rate was consistent across groups at the same foraging stage (GLMM: χ² = 0.77, df = 2, *p* = 0.68). We analysed the foraging behaviour of each bee group in more detail:

#### (A) Low pickiness bees

Low pickiness bees maintained consistent foraging durations across bouts (post-hoc Tukey test, bout 1 vs. bout 3: 3.89 min ± 0.34 vs. 3.17 min ± 0.45, z-ratio = 1.01, *p* = 0.57; Fig. 3A). They increased the number of flowers collected (post-hoc Tukey test, bout 1 vs. bout 3: 10.8 flowers ± 1.02 vs. 12.8 flowers ± 1.18, z-ratio = −2.70, *p* = 0.019; Fig. 3B) but did not significantly increase the proportion of high-concentration sucrose solution collected (post-hoc Tukey test, bout 1 vs. bout 3: 39.1% of high-quality flowers collected ± 6.74 vs. 55.2% ± 6.57, z-ratio = −1.81, *p* = 0.17; Fig. 3C).

**Figure 3:**
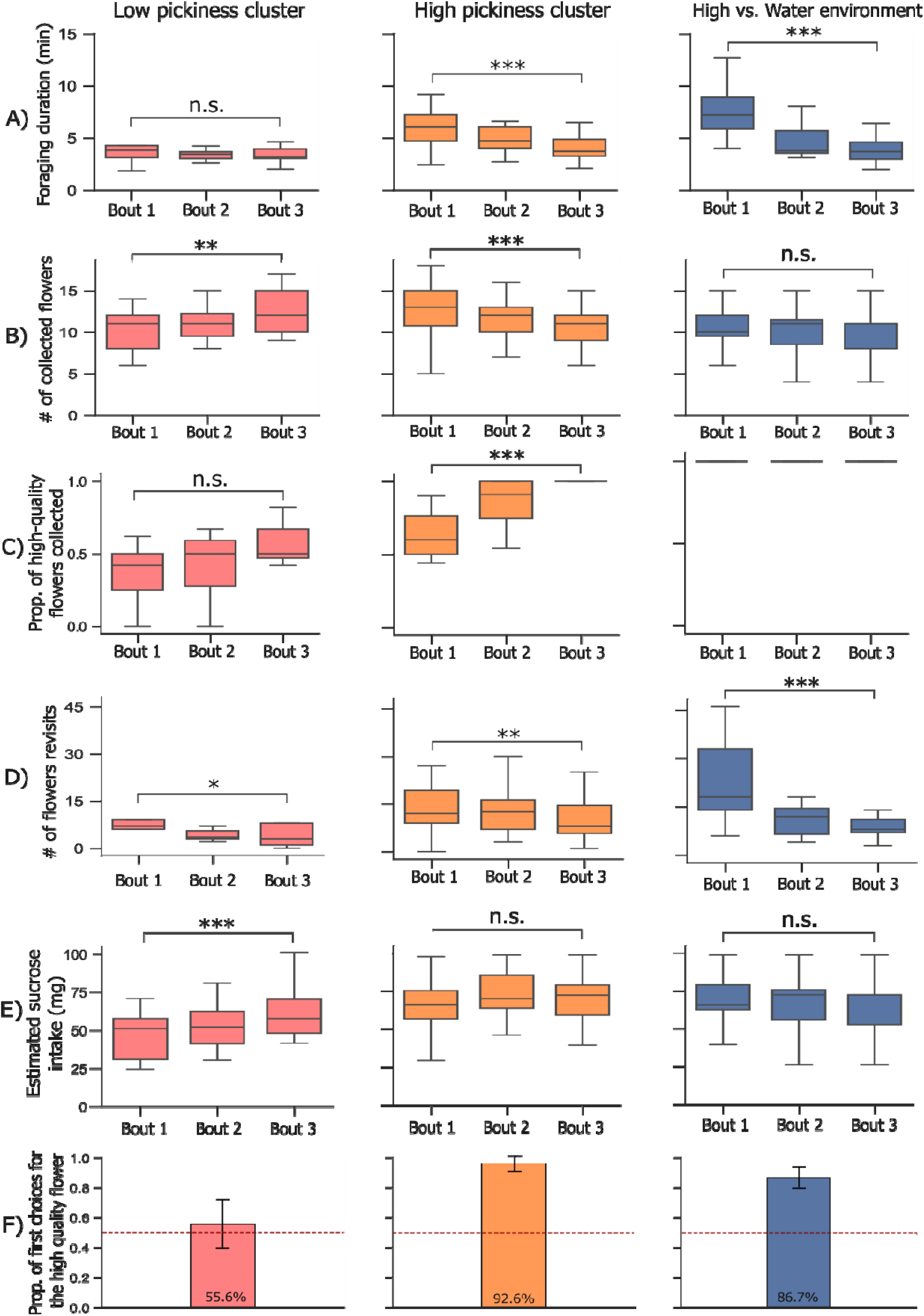
Foraging behaviour across environments and pickiness levels. **A)** Foraging duration per foraging bout (minutes); **B)** Number of flowers collected per bout; **C)** Proportion of high-concentration sucrose solution collected per bout; **D)** Number of flowers revisited per bout; **E)** Estimated sucrose intake per foraging bout (mg), and **F)** Proportion of bees whose first visit was to the high-quality flower in the final test (N = 52 bees). Each boxplot shows the median (centre line), the interquartile range (IQR), and whiskers extending to the smallest and largest values within 1.5×IQR. Error bars represent standard errors of the predicted probabilities. In panel C, probabilities for the High vs. Water environment are equal to 1 in each bout because only high-quality flowers were available. Statistical significance is indicated as **\*\*\*** (*p* < 0.001), **\*\*** (*p* < 0.01), **\*** (*p* < 0.05), and **n.s.** (*p* > 0.05, post-hoc Tukey tests).

The bees reduced revisits to previously depleted flowers (post-hoc Tukey test, bout 1 vs. bout 3: 7.57 revisits ± 1.87 vs. 4.7 revisits ± 1.23, z-ratio = 2.63, *p* = 0.02; Fig. 3D). Overall, bees from this cluster showed an increase in estimated sucrose intake across bouts (post-hoc Tukey test, bout 1 vs. bout 3: 48.1 mg ± 5.07 vs. 64.2 mg ± 6.58, z-ratio = −4.47, *p* < 0.0001; Fig. 3E). In the final test, 55.6% of bees first visited the high-quality flower patch (Fig. 3F).

#### (B) High pickiness bees

By contrast, High pickiness bees decreased their foraging duration over bouts (post-hoc Tukey test, bout 1 vs. bout 3: 6.39 min ± 0.56 vs. 3.89 min ± 0.34, z-ratio = 6.18, *p* < 0.0001; Fig. 3A). They decreased the number of flowers collected (post-hoc Tukey test, bout 1 vs. bout 3: 13.2 flowers ± 0.76 vs. 11 flowers ± 0.64, z-ratio = 5.12, *p* < 0.0001; Fig. 3B) while strongly shifting their collection of sucrose solution towards high-quality flowers (post-hoc Tukey test, bout 1 vs. bout 3: 61.6% of high-quality flowers collected ± 3.59 vs. 97.6% ± 1.01, z-ratio = −7.18, *p* < 0.0001; Fig. 3C).

The bees reduced flower revisits (post-hoc Tukey test, bout 1 vs. bout 3: 12.73 revisits ± 1.99 vs. 9.88 revisits ± 1.57, z-ratio = 3.26, *p* = 0.003; Fig. 3D), while their estimated sucrose intakes remained stable across bouts (post-hoc Tukey test, bout 1 vs. bout 3: 68 mg ± 4.27 vs. 71.8 mg ± 4.5, z-ratio = −1.69, *p* = 0.21; Fig. 3E). In the final test, 92.6% of bees first visited the high-quality flower patch (Fig. 3F).

#### (C) Bees in the High vs. Water flower environment

In this flower environment, where only high-concentration sucrose solution was available, bees nearly halved their foraging duration over successive bouts (post-hoc Tukey test, bout 1 vs. bout 3: 6.74 min ± 0.83 vs. 3.89 min ± 0.34, z-ratio = 5.96, *p* < 0.0001; Fig. 3A) while maintaining a steady number of flowers collected (post-hoc Tukey test, bout 1 vs. bout 3: 10.8 flowers ± 0.91 vs. 10.1 flowers ± 0.86, z-ratio = 1.25, *p* = 0.42; Fig. 3B).

Bees reduced flower revisits by nearly threefold (post-hoc Tukey test, bout 1 vs. bout 3: 24.6 revisits ± 4.95 vs. 9.01 revisits ± 1.9, z-ratio = 10.64, *p* < 0.0001; Fig. 3D), while their estimated sucrose intakes remained consistent across bouts (post-hoc Tukey test, bout 1 vs. bout 3: 71.5 mg ± 6.26 vs. 66.5 mg ± 5.85, z-ratio = 1.61, *p* = 0.24; Fig. 3E). In the final test, 86.7% of bees first visited the high-quality flower patch (Fig. 3F).

## Discussion

This study investigated how differences between high- and low-quality flowers in mixed-quality patches influence bees’ flower choices, addressing a gap in our understanding of interindividual variability in bumblebee foraging behaviour. We found that foragers rapidly improved their efficiency across successive bouts, reaching similar sucrose intake rates by the third bout regardless of flower environment type, whether the environment was “favourable” (containing both high- and low-quality flowers) or “challenging” (where only half the flowers were rewarding). This finding supports our prediction and aligns with Optimal Foraging Theory (OFT), which posits that nectarivores should maximise their net energy intake rate (Heinrich, 1979a; Waddington and Holden, 1979; Pyke and Starr, 2021). Additionally, we observed substantial interindividual variability in foraging behaviour, particularly in flower selectivity: some bees strongly preferred high-quality flowers, while others were less selective. This is consistent with previous findings of high individual variability in bumblebee foraging behaviour in heterogeneous environments (Saleh and Chittka, 2007; Lihoreau et al., 2011; Woodgate et al., 2016).

A key prediction of our study was that bees in the challenging environment would more quickly learn to distinguish high-quality flowers, due to the stronger contrast between flower types. This was supported by their increased visits to high-quality flowers in foraging bouts 2 and 3, compared to bees in the favourable environment. While this may reflect faster learning, bees in the favourable environment may have intentionally continued visiting and collecting both flower types, as both offered some reward (see next section). Notably, bees in the challenging environment were even more selective than high-pickiness bees in the favourable environment, suggesting that harsher conditions may have indeed accelerated learning. Nevertheless, bees in both environments ultimately achieved similar estimated sucrose intake rates in each bout, highlighting their behavioural flexibility in adapting to different environments.

Bees in the favourable environment showed marked individual differences in how they improved foraging efficiency. High-pickiness bees collected smaller volumes of more concentrated sucrose, prioritising quality over quantity. In contrast, low-pickiness bees increased the total volume collected, regardless of concentration. Although both groups learned and improved over successive bouts, these differences in foraging behaviours remained consistent over time. This mirrors findings by Chittka et al. (2003), where individual bees consistently differed in speed-accuracy trade-offs throughout learning. Similar behavioural trade-offs have been reported in other studies, with bees balancing speed and accuracy (Burns and Dyer, 2008) or balancing reward value against foraging effort (Lihoreau et al., 2011; Pattrick et al., 2023). Such individual variability may be adaptive in unpredictable environments, where resource quality and availability vary across space and time. Moreover, it can enhance colony performance by diversifying foraging roles (Muller and Chittka, 2008; Holland et al., 2021). Supporting this, Hagbery and Nieh (2012) found that generalist bumblebees, those not specialising strictly in nectar or pollen, adjusted their foraging behaviour flexibly in response to changes in colony needs.

Interestingly, most bees did not increase their estimated absolute sucrose intake across bouts; only low-pickiness bees showed an increase. Bees in this cluster had lower estimated intakes during their first two foraging bouts compared to both high-pickiness bees and bees foraging in the challenging environment with only one rewarding flower type. For high-pickiness bees, their focus on high-quality flowers likely constrained their intake over time. In our setup, only half the flowers in each patch were rewarding, increasing the likelihood of revisiting depleted flowers as the best rewards were collected. Previous studies have shown that bees do not visit all flowers in a patch (Goulson, 2000; Hemingway et al., 2024), and according to the optimal departure rule, pollinators should leave a patch when their reward intake rate declines (Cresswell, 1990; Goulson, 2000).

However, we found that naïve bees, regardless of group, did not collect substantially less sucrose during their first foraging bout, despite longer foraging times and frequent revisits to empty flowers. This suggests that bees did not abandon patches based on declining reward rates; if they had followed an optimal departure rule, we would expect lower sucrose intakes in early bouts. That said, our setup offered no alternative foraging sites, so bees may have continued exploring simply because there was nowhere else to go. Their behaviour likely reflects exploration of a novel environment, similar to the orientation flights seen in wild bumblebees (Woodgate et al., 2016).

Surprisingly, high-pickiness bees in the favourable environment collected over 15% fewer flowers in their final bout compared to their first, while maintaining stable sucrose intake. This pattern suggests that bees may return to the nest after reaching a fixed energetic threshold, rather than filling their crop or foraging for a set duration. It also supports the idea that bees may return with partially empty crops (see Schmid-Hempel et al., 1985). However, explicit testing is necessary to confirm this fixed energetic target hypothesis.

One result worth further consideration is that all bee groups in our study improved their foraging efficiency at a similar rate, reaching comparable estimated sucrose intake rates at each bout. This suggests that bees may be capable of compensating for less favourable initial conditions by adjusting their foraging behaviour. However, this finding should be interpreted with caution, as we did not directly measure sucrose intake but instead inferred it through video analysis and calculations (refer to Data collection and analysis section). Importantly, our estimates may underestimate intake from low-quality flowers, which were often probed for less than a second and thus classified as non-collected. Yet, Lechantre et al. (2021) demonstrated that bumblebees can imbibe up to 2 µL of solution per second, suggesting that even very brief visits may still result in meaningful reward intake.

Our study also raises questions about what shapes individual foraging behaviours, especially flower-quality pickiness. In the favourable environment of high- and low-quality flowers, most bees quickly became highly selective for the high-quality flowers. Notably, pickiness was associated with bees’ initial flower choices, which were most often to high-concentration sucrose flowers. While this pattern may reflect reward-driven learning, it is also possible that innate colour preferences influenced flower choices, as most high-concentration flowers in the main experiment were yellow. To test this, we reversed the colour-reward associations in a complementary experiment and included experiment type (main vs. complementary: yellow vs. blue as the high-quality flower) as a fixed effect in all statistical models. This variable had no significant effect on foraging behaviours, suggesting that the observed pickiness was primarily driven by reward quality rather than colour.

Previous studies have shown that bees tend to remain constant to flower types offering high rewards but explore alternative options when rewards are low (Chittka et al., 1999; Grüter and Ratnieks, 2011; Nityananda and Chittka, 2020). This could explain why bees in our study strongly favoured high-quality flowers after initial exposure, whereas those first encountering low-quality flowers displayed more variability in their flower choices. Although flower constancy was long considered a cognitive limitation (Waser, 1986; Chittka et al., 1999), more recent work suggests it can function as an adaptive strategy (Gegear and Thomson, 2004). Supporting this, recent model simulations have shown that colonies composed of highly flower-constant individuals exploited only 30–50% of available flower species, whereas colonies of indiscriminate foragers could exploit nearly all species (Grüter et al., 2024).

Can pickiness be solely attributed to the quality of a bee’s first few flower visits? The moderate strength of the correlation suggests that other factors likely influenced pickiness as well. For instance, bees vary in their sucrose sensitivity thresholds, which may drive differences in preference for reward quality (Page et al., 2006; Riveros & Gronenberg, 2010). Additionally, differences in learning ability among bumblebees (Raine et al., 2006) might explain why bees in the low pickiness cluster did not favour the high-quality patch in the final test. While all bees in our study increased their visits to high-quality flowers over successive bouts, they did so at different rates. Would every bee have eventually become highly selective in our setup? Adjusting foraging behaviour can be energetically costly, especially in dynamic environments (DeWitt et al., 1998). A promising research direction would be to manipulate initial conditions to test whether, and when, bees converge on the same optimal foraging behaviour. This could provide insight into the limits of behavioural flexibility.

It is worth noting that most high-concentration flowers in our experiment were yellow. We had expected bees to favour blue flowers on their first visit, given well-documented innate preferences (Gumbert, 2000; Raine & Chittka, 2007). However, over two-thirds of initial visits were to yellow flowers. This unexpected bias may be due to yellow flowers reflecting more light across a broader wavelength range, making them appear brighter overall (see reflectance curves in **S3**). Brightness is known to strongly influence bee preferences (Sletvold et al., 2016) and may explain this initial preference, even though blue flowers in our setup had higher spectral purity and saliency in the white arena—traits typically associated with greater attractiveness to bees (Lunau, 1990; Goulson, 2000).

While our findings suggest that bees aimed to optimise energy intake, not all energetic costs of foraging were accounted for. For example, flight costs were minimal in our setup, as flowers within patches were close enough for bees to walk between them, potentially conserving energy. Energy expenditure also likely varied with the weight of the sucrose loads carried by each bee (Wolf et al., 1989; Combes et al., 2020). Although bees learned quickly in our simple flower environments over just a few bouts, natural environments are far more complex, with less predictable and more subtle differences between high- and low-reward flowers. Our small flight arena also represented a “local” foraging scale (Sommer et al., 2022), where bees likely encountered floral cues almost immediately (Heinrich, 2004)—a contrast to natural foraging conditions. Still, our findings highlight the rapid adaptability of bumblebee foraging behaviour and point to promising future research directions: exploring how environmental conditions, initial experience, and foraging behaviour interact could provide new insights into the ecological success of bumblebees across diverse and dynamic habitats.

## Supplementary Material

**S1** presents the statistical analysis of the main experiment, while **S2** provides the statistical analysis of the complementary experiment. **S3** includes the reflectance curves of the artificial flowers, and **S4** contains the reflectance dataset. **S5** shows the layout patterns of the flower environments. **S6** features video examples of bees foraging during the experiment. **S7** and **S8** contain the datasets for the main and complementary experiments, respectively. All statistical analyses and datasets are publicly available on Zenodo (https://doi.org/10.5281/zenodo.16418595).

## Acknowledgments

We thank Dorian Champelovier for his help with data collection, and the anonymous reviewers for their thoughtful feedback which helped improve the manuscript.

